# BetaAlign: a deep learning approach for multiple sequence alignment

**DOI:** 10.1101/2024.03.24.586462

**Authors:** Edo Dotan, Elya Wygoda, Noa Ecker, Michael Alburquerque, Oren Avram, Yonatan Belinkov, Tal Pupko

## Abstract

The multiple sequence alignment (MSA) problem is a fundamental pillar in bioinformatics, comparative genomics, and phylogenetics. Here we characterize and improve BetaAlign, the first deep learning aligner, which substantially deviates from conventional algorithms of alignment computation. BetaAlign draws on natural language processing (NLP) techniques and trains transformers to map a set of unaligned biological sequences to an MSA. We show that our approach is highly accurate, comparable and sometimes better than state-of-the-art alignment tools. We characterize the performance of BetaAlign and the effect of various aspects on accuracy; for example, the size of the training data, the effect of different transformer architectures, and the effect of learning on a subspace of indel-model parameters (subspace learning). We also introduce a new technique that leads to improved performance compared to our previous approach. Our findings further uncover the potential of NLP-based approaches for sequence alignment, highlighting that AI-based methodologies can substantially challenge classic tasks in phylogenomics and bioinformatics.

## Introduction

The Needleman–Wunsch algorithm was the first to use dynamic programming to efficiently find the best global scoring alignment between two sequences (Needleman & Wunsch, 1970). The inference of a multiple sequence alignment (MSA) was later shown to be an NP-complete problem (Wang & Jiang, 1994), making the inference task impractical for a large set of sequences. To overcome this hurdle, popular MSA algorithms, such as MAFFT (Katoh & Standley, 2013) and PRANK (Löytynoja, 2014), use heuristics to reduce the search space and consequently, the running time.

There is extensive knowledge regarding the variability of the evolutionary process among different datasets and lineages. For example, amino-acid replacement matrices vary between proteins encoded in the nuclear genome, the mitochondria, and plastids (Pesole et al., 1999). Indel dynamics also highly vary between datasets and among different phylogenetic groups (Ajawatanawong & Baldauf, 2013; Loewenthal et al., 2021; Wolf et al., 2007). Furthermore, site-specific evolutionary rates vary along the analyzed sequence. For example, amino-acid sites that are exposed to the solvent tend to have higher evolutionary rates compared to buried sites (Wang et al., 2008). Alignment algorithms using default configurations implicitly assume that the evolutionary dynamics do not vary among different datasets and within a single dataset. The general inability of MSA inference algorithms to automatically tune their scoring scheme to the specific dataset being analyzed is a shortcoming of present alignment programs. The “one matrix fits all biological datasets” and “one matrix fits all regions within a dataset” assumptions implicitly employed by current methodologies raise fundamental questions about the correctness of alignments produced by such methods. Although it is possible to modify gap-penalty parameters in some alignment program, these programs do not provide means to automatically tune the parameters to specific datasets or regions within a dataset, and hence, by and large, all users employ the default settings.

Alignment algorithms are typically assessed by empirical alignment regions, but these regions are not comprehensive enough to cover the entire range of alignment challenges. It is worth noting that these regions are often calculated manually, so their reliability as a “gold standard” is uncertain (Iantorno et al., 2014). Many differences may exist between empirical and simulated datasets, e.g., the former may include evolutionary scenarios that are not modeled in simulations such as micro-rearrangements (Walker et al., 2021). Thus, when alignment programs are tested with simulated complex alignments, the results often substantially differ from empirical benchmark outcomes (Chang et al., 2014).

One of the key concepts in learning algorithms, in general, and in deep-learning algorithms in particular, is the ability to learn from previously annotated data, i.e., to generalize from previous observations to unseen cases. For the task of alignment inference, a deep-learning algorithm should learn from “true” alignments (e.g., simulated sequences for which the correct alignment is known) and apply the obtained knowledge to align novel sequences. In this work, we aimed to harness natural language processing (NLP) learning algorithms to the task of aligning sequences, thus to better capture the evolutionary dynamics of biological sequences.

Here we present an evidently effective improvement for our previously developed BetaAlign approach (Dotan, et al., 2023a). Instead of computing a single alignment, we now calculate multiple alternative alignments and return the alignment that maximizes the certainty, leading to significant performance improvement. To further characterize BetaAlign, we conducted the following analyses: (1) evaluating the effect of training time and size; (2) measuring the performance as a function of the evolutionary dynamics that generated the sequences; (3) evaluating the effect of transfer learning; and (4) comparing different transformer architectures. We also introduce the term subspace learning to describe training on a subspace of the indel parameters and investigate its utility for BetaAlign. Lastly, we show that the benefit of our approach is also transferable, that is, the embedding obtained by the model could serve as meaningful features for accurate inference in other learning tasks such as root length prediction. Table 1 describes the main differences between the previous and current work. For completeness, we start by describing the algorithm.

**Table 1.**
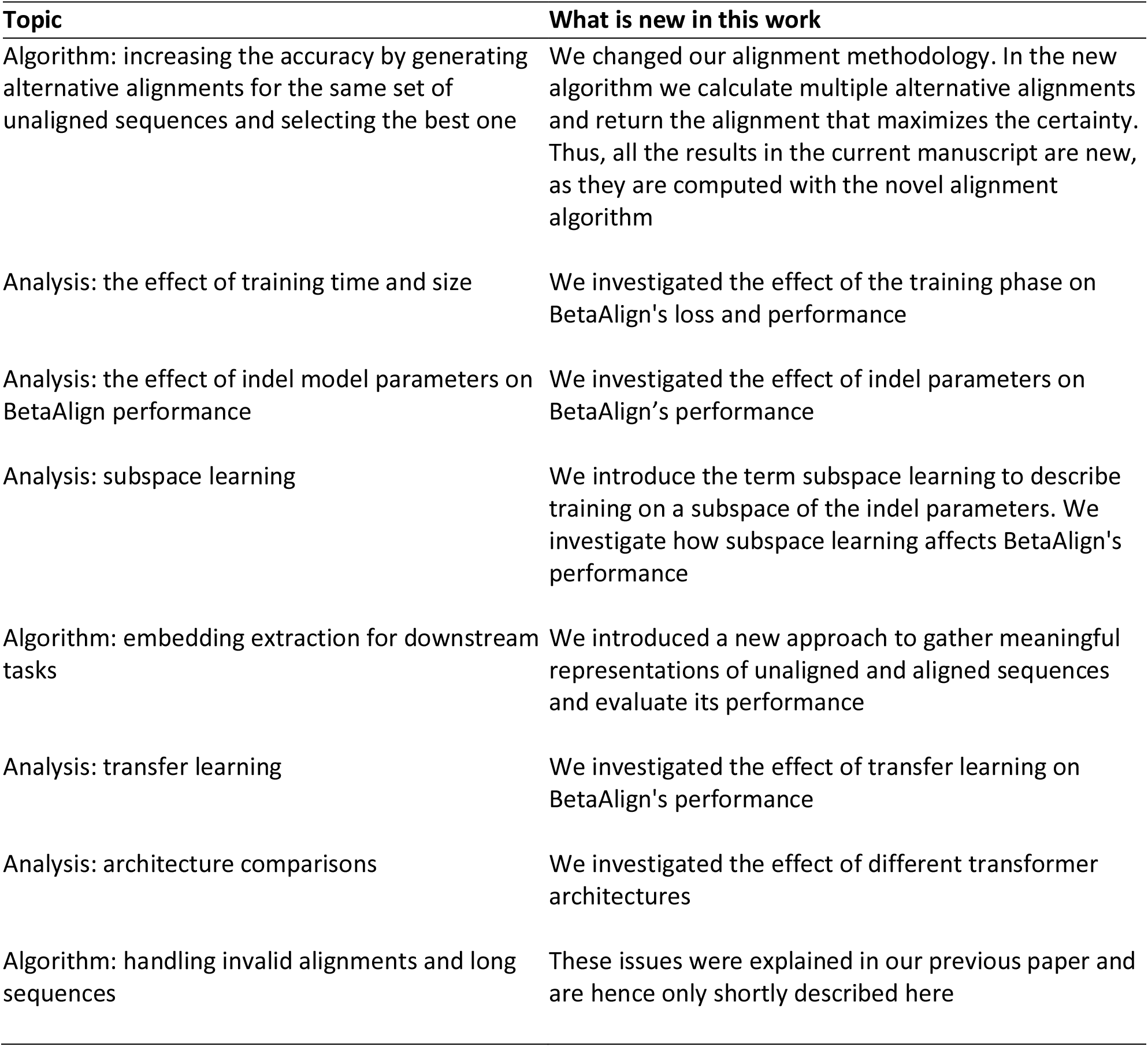
The different topics discussed in this research compared to the previous version of BetaAlign.

## New Approach

### Outline

Typically, sequence-to-sequence NLP tasks involve a single sentence (or text) as both input and output, e.g., translating from one language to another or changing a sentence from active to passive (Bahdanau et al., 2016; Shalumov & Haskey, 2023; Sutskever et al., 2014). The learning phase of the algorithm is to map a single input sentence to a single output sentence. When we aim to apply sequence-to-sequence models to the problem of alignment, we are faced with a problem: the input to the alignment task is several “sentences”, each corresponding to an unaligned sequence. Similarly, the output is a set of related sentences, each corresponding to a row in the resulting alignment. The first task in the BetaAlign algorithm is to transform the set of unaligned sequences to a single “sentence”. Such input-transformation can be done, e.g., by concatenating all the unaligned sequences, adding a special character (we use the pipe character, “|”) to indicate the boundaries between the sequences (Fig. 1). For training the algorithm, we also need to provide target sentences. Thus, we also need an output-transformation step, in which we convert resulting alignments to a single target sentence. In BetaAlign we use the “*spaces*” representation (Fig. 1). The above representations allow providing a sequence-to-sequence model with a large set of examples of valid source and target sentences, which are used for model training. The models that we use rely on the transformer architecture (Vaswani et al., 2017). Once trained, the optimized transformer can process new unseen examples, in our case, it can transform (unseen) unaligned sequences to an alignment.

**Fig. 1.**
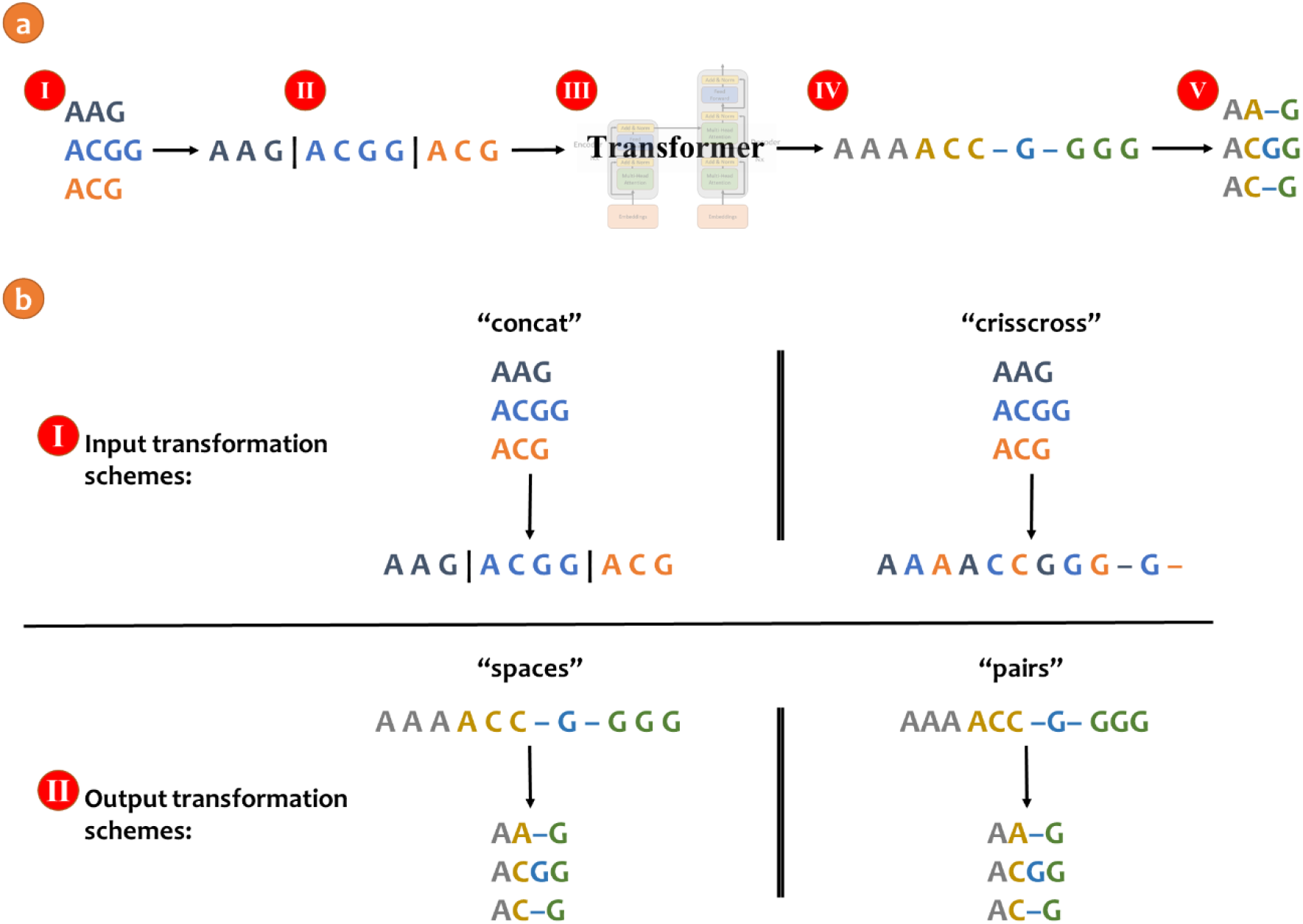
Example of aligning three sequences with BetaAlign (a): (Ⅰ) Consider the unaligned sequences “AAG”, “ACGG” and “ACG”; (Ⅱ) The unaligned sequences are concatenated to a single sentence with a special character “|” between each original sequence; (Ⅲ) The trained model processes the single input sentence and generates the single output sentence; (Ⅳ) The processed output is structured such that the first three nucleotides represent the first column, the next three nucleotides represent the second column, and so on; (Ⅴ) The output is converted into an MSA. (b) An illustration of the different input (Ⅰ) and output (Ⅱ) transformation schemes.

There are several aspects that need to be addressed to fully describe the BetaAlign algorithm and how its performance was evaluated. These include, for example, the generation of training and test data, the transformer architecture and how it was trained, the handling of long sequences and how the generation of invalid alignments was prevented. We aim to provide a more general description as part of the New Approach section, while technical details are provided in the Methods section.

### The generation of training and test data

For both training and testing the performance of BetaAlign, many sets (data points) of unaligned sequences and their corresponding “true alignment” were needed. These data points were generated using simulations. Specifically, we use SpartaABC (Loewenthal et al., 2021), which allows different length distributions for insertions and deletions. For example, the initial testing and training for the pairwise alignment problem were achieved by generating millions of pairs of two unaligned sequences and their corresponding MSAs for the training and testing data. The indel rates, their type (insertion or deletion), and their length distribution were sampled from specific ranges. We note that we do not assume equal rates of insertions and deletions, nor equal length distributions for the two types of events (this is mainly important when simulating along a tree rather than when simulating pairwise alignments).

### The transformer architecture

Transformers are currently the working horse of NLP and other AI domains. A transformer is a deep-learning model designed to handle discrete sequential data. The transformer used in our work is composed of an encoder and a decoder. The encoder embeds each input sequence into a sequence of high-dimensional vector representation. Next, the decoder receives those representations and the last generated word and predicts the next word. Transformers may vary in architecture, number of layers, and size, which corresponds to the number of tunable architectural hyper-parameters. When training a transformer, one can also vary the learning hyper-parameters, e.g., the parameter “max tokens” determines how much input to process before the model parameters are updated. We have tested several transformer architectures and parameters, implemented using the Fairseq library (Ott et al., 2019). Technical details regarding transformer optimizations are provided in the Methods.

### Transfer learning and subspace learning

The input and output patterns of the analyzed sequences vary as a function of their number, e.g., the number of pipe characters in the “*concat*” representation. We thus optimized a different transformer for each number of sequences. To this end, when optimizing the transformer for, say, five sequences, we start the parameter optimization step from the set of optimal parameters obtained for the previous transformer that was trained on four sequences, a technique called transfer learning (Avram et al., 2023; Tan et al., 2018).

We also use transfer learning in order to train a transformer on sub-regions of the parameter space, i.e., subspace learning (see Methods). For example, we can train a general pairwise alignment transformer as described above and then train a different transformer only for alignments with a high ratio of indels to substitutions. In essence, this allows training several transformers, specialized for sub-regions of the parameter space.

### Handling invalid alignments

Transformers have no inherent mechanism that restricts them to generate valid alignments. Thus, in some cases, a trained transformer may produce invalid output. For example, when aligning sequences, each output sequence, including gap characters, should have the same length (Fig. 2). To this end, we trained several different transformers, which differ from each other with respect to their tunable hyper-parameters, on the same training dataset (see Methods). If a transformer provided an invalid alignment, we provided the output of an alternative transformer.

**Fig. 2.**
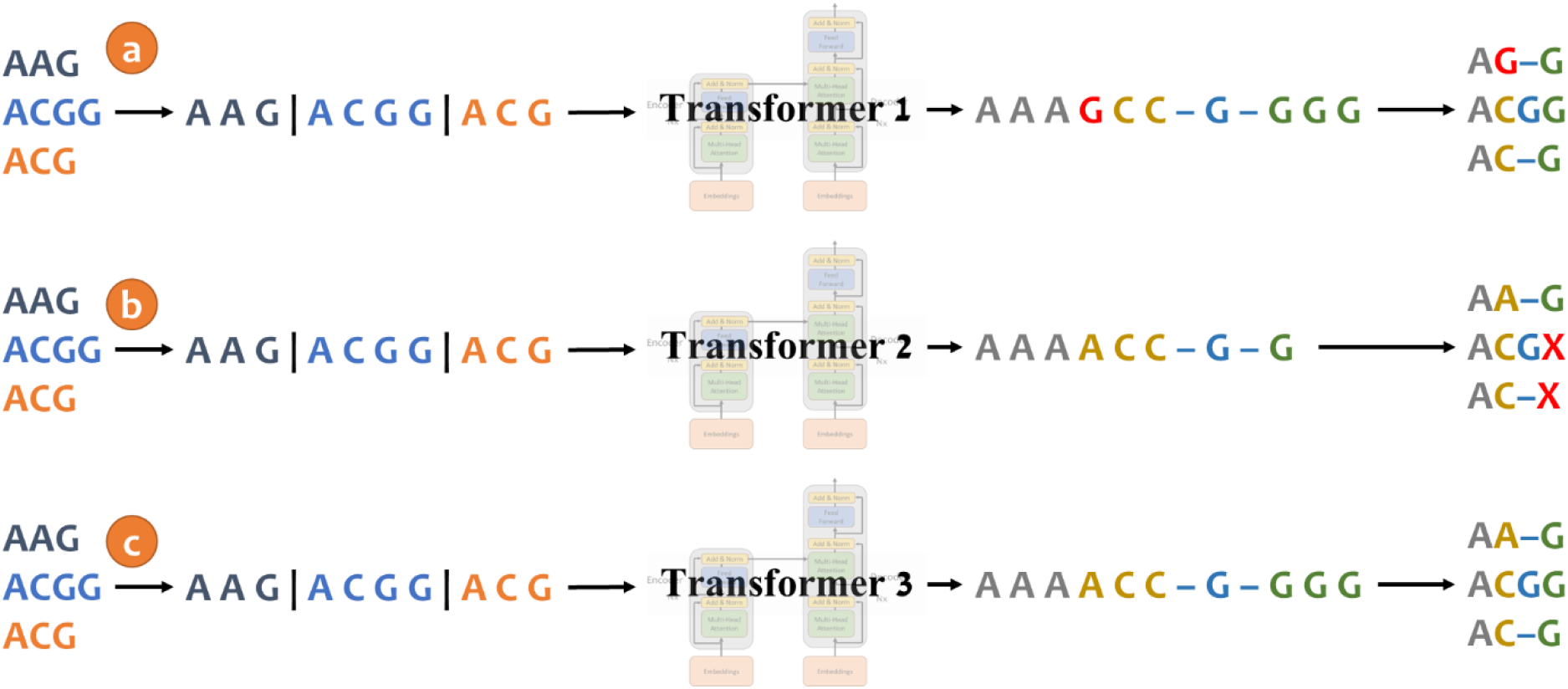
Example of handling invalid alignments. Consider the unaligned sequences from Fig. 1: “AAG”, “ACGG” and “ACG”. (a) When aligning these sequences, BetaAlign mistakenly mutated the character “A” to “G” (red); (b) Aligning the same sequences with a different transformer resulted in a different output, but here the transformer generated a shorter sequence in which the last two characters are missing (the red “X” was added to mark the missing nucleotides); (c) The third transformer provided a valid alignment as output and can be used as the output of BetalAlign.

### Handling long sequences

The transformers that we have utilized were designed to process text of natural languages and not biological sequences. As such, they are limited to processing sentences with up to 1,024 tokens (a token in natural languages is the building block of a sentence, in our case, each token is either a base pair or an amino acid). When aligning biological sequences, the input and output sentences often exceed this length threshold. Due to memory and run-time constraints, increasing the threshold is infeasible. To overcome this challenge, we introduced a “segmentation” methodology, in which we align segments of the alignments, which are later concatenated to form the entire MSA. This procedure is achieved by training dedicated transformers for this task (Dotan, et al., 2023a).

### Considering alternative input and output transformation schemes

The transformer architectures we harnessed for the task of aligning sequences are sequence-to-sequence models. One of the key components of our proposed alignment approach is to transpose the multiple input sequences into a single sentence that can be processed by the transformer. Input transformation converts the unaligned sequences into the “input sentence” of the transformer while output transformation converts the “output sentence” of the transformer into an MSA.

There are various transformation schemes available for converting unaligned sequences into a single sentence. In Fig. 1a we presented the “*concat*” representation: the unaligned sequences are concatenated with a special character “|”. The vocabulary, which encompasses the entire set of possible tokens, of this scheme is {“A”, “C”, “G”, “T” and “|”} for the nucleotide sequences. We used the “*spaces*” representation for output transformation, in which each of the amino acids or nucleotides is considered a separate token. The vocabulary of this scheme is {“A”, “C”, “G”, “T” and “–”} for the nucleotide sequences.

However, alternative transformation schemes for the source sequences can be considered. We previously considered the “*crisscross*” scheme, the tokens of the unaligned sequences are interleaved (Dotan, et al., 2023a). That is, the first token represents the first character from the first unaligned sequence, the second token represents the first token of the second unaligned sequence, and so on. The vocabulary of this scheme is {“A”, “C”, “G”, “T” and “–”} for the nucleotide sequences. Of note, the gap character is used to fill the gaps if the sequences are of different lengths (Fig. 1b). Similarly, alternative transformation schemes for generating the output sentence are possible.

In the “*pairs*” scheme each token represents the entire column. The vocabulary of this scheme depends on the number of unaligned sequences, for instance, when aligning three DNA sequences the vocabulary size is 124 tokens: {“AAA”, “AAC”, “AAG”, “AAT”, “AA–”,…, “TTG”, and “TTT”}. Of note, the token “– – –” (three gap characters) is invalid as such column cannot exist.

It is important to remember that the transformation schemes are external to the transformer itself. Each transformation methodology creates a different mapping from unaligned sequences to an MSA, which requires training the transformer on these representations. Different considerations come into play when selecting the appropriate scheme (Dotan, et al., 2023a). In the “*pairs*” scheme, the output sequence length is the number of columns while in the “*spaces*” the length is the number of nucleotides. Because length is a limiting factor when using current transformer architectures, using the “*pairs*” scheme may be advantageous. However, the “*pairs*” scheme restricts the use of transfer learning (see below). When transitioning from pairwise alignment to aligning three sequences, the vocabulary would change (from 24 tokens to 124 tokens) and in general, the number of possible tokens exponentially increases as a function of the number of unaligned sequences. In our previous work we observed that the “*concat*” and “*spaces*” representations (shown in Fig. 1a) performed best (Dotan, et al., 2023a). Thus, all the experiments in this work are done with these representations for the input sequences and output MSA, respectively.

### Increasing the accuracy by generating alternative alignments for the same set of unaligned sequences and selecting the best one

We present a method for generating multiple alternative MSAs from the same input data. This is done by randomizing the order in which the input unaligned sequences are concatenated (see Methods). We also show how we select a single MSA from this set using a “majority voting” approach. We show that this data augmentation followed by majority voting approach provides a more accurate MSA than relying on a randomly sampled MSA from the set of alternative MSAs, on average. The majority voting approach relies on computing for each MSA, the degree of its agreement with all other alternative MSAs and selecting the one that agrees the most (see Methods).

## Results

### The effect of training time and size

We tested how the number of epochs (a single pass on the whole training set) and training size affect the accuracy and coverage of BetaAlign. We compared the model’s performance when trained on three training data sizes: 50,000, 100,000 and 200,000 protein alignments. Our results clearly indicate that for all datasets, the training loss decreases as the number of epochs increases, reaching almost a plateau when the data size is 200,000 alignments (Fig. 3). For each training data size, the validation loss follows the decrease in the training loss, suggesting that there is no over-fitting for the transformer. The coverage (fraction of resulting alignments that are valid) also continuously increases, e.g., after 20 epochs the coverage was ∼ 40% while after 60 epochs, the coverage was already ∼80%.

**Fig. 3.**
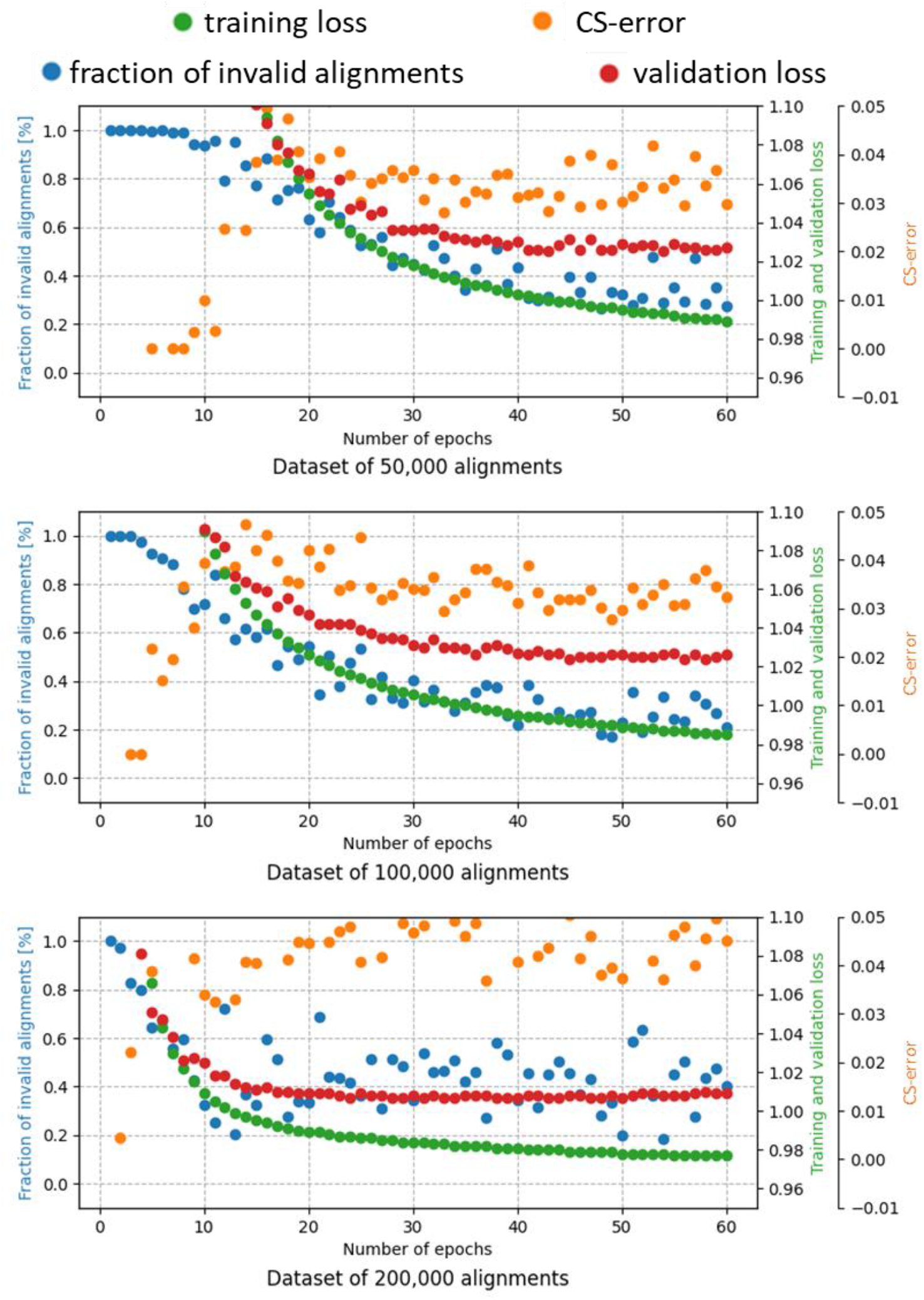
Effect of increasing the training time (number of epochs) and size (number of different MSAs) on the fraction of invalid alignments (blue dots), CS-error (orange dots), validation loss (red dots), and training loss (green dots). All alignments were of three protein sequences, dataset SPD2. Note that the figure contains the four metrics together for comparing the correlation between the metrics. Each metric has a different range, and thus, there are multiple y-axes. Also note that the errors and coverage in this graph are based on a single alternative alignment, while in practice both the accuracy and coverage are substantially improved by considering a set of alternative MSAs (see text for details).

The CS-error seems to substantially fluctuate even after 30 epochs (we note that the CS-error quantifies the error on valid alignments only, while the loss function quantifies the error on all alignments). The results further suggest that the loss function is correlated to the CS-error, but the correlation is mediocre at best. The correlation between the loss on the validation data and the CS-error on the dataset of 100,000 alignments, between epochs 20 and 60 was *R*^2^ = 0.467 (*P* = 0.0023). We speculate that this low correlation reflects the fact that the loss function is different from the CS-error.

Comparing the training and validation loss between the different training size datasets indicated that increasing the training size decreases the loss as expected (training loss at epoch 60: 0.989, 0.985, 0.977, for datasets of 50,000, 100,000, 200,000, respectively). This gain in accuracy as reflected in the loss function was not evident when the performance is measured by the CS-error, probably reflecting the mediocre correlation between the two scores discussed above.

### The effect of indel model parameters on BetaAlign performance

We next studied the effect of the different indel parameters (of the assumed indel model that generated the simulated data) on the performance. To this end, we divided the alignments into bins by their evolutionary parameters: the insertion and deletion rate parameters (*R*_*I*_ and *R*_*D*_, respectively) and the parameters that determine the distribution of indel lengths (*A*_*I*_ and *A*_*D*_ for the insertion and deletion distributions, respectively). As expected, increasing the indel rate parameters *R*_*I*_ and *R*_*D*_ substantially decreases accuracy (Fig. 4a). The size distribution of the indels had little effect on accuracy (Fig. 4b).

**Fig. 4.**
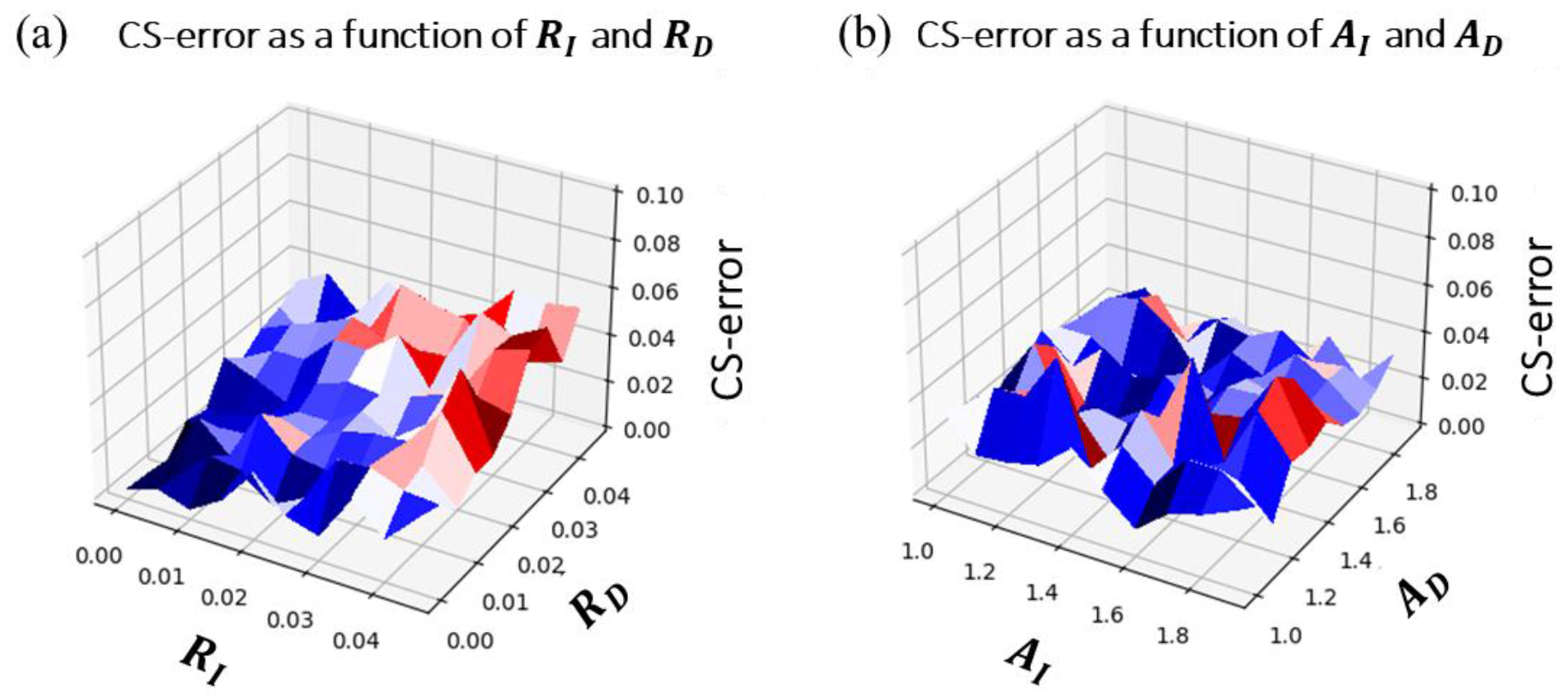
The effect of indel parameters on BetaAlign performance: (a) The effect of *RI* and *RD* (in this case *AI* and *AD* were sampled from the entire range); (b) The effect of *AI* and *AD* (in this case *RI* and *RD* were sampled from the entire range). Figure illustrates the results on protein dataset SPD3.

### Subspace learning

As stated above, we can train a transformer on a set of MSAs that share specific features, e.g., training them on MSAs with a high deletion rate and a low insertion rate. To determine if such a subspace-learning approach increases accuracy, we simulated three nucleotide datasets of five sequences per sample (see Methods). The first dataset, “general” (ND10), was simulated with a wide range of indel model parameters. The second dataset, “specific” (ND11), was simulated on a sub-space of the indel model parameter space, i.e., the generated MSAs resemble each other in terms of indel dynamics. Finally, the third dataset, “ultra-specific” (ND12), is even more restrictive in terms of the allowed indel dynamics (see Table S2). Our results suggest that subspace learning can improve both coverage and accuracy (Fig. 5), with a more substantial effect on coverage. This highlights the importance of fitting the correct configuration of the alignment program (and in our case the training of the transformer) to the specific data. These results demonstrate that subspace learning has the potential to improve the accuracy of BetaAlign.

**Fig. 5.**
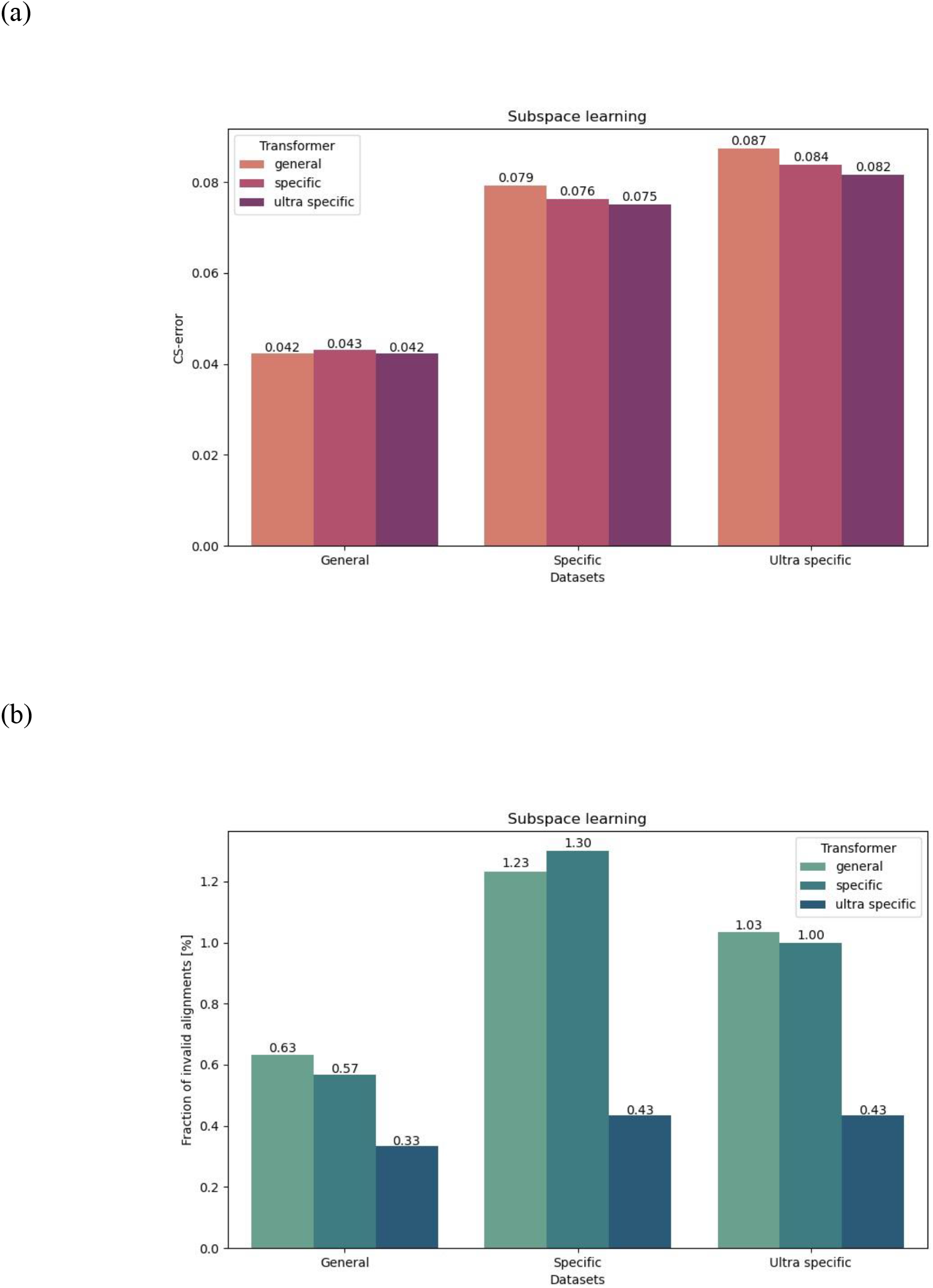
Effect of subspace learning on the CS-error (a) and the fraction of invalid alignments (b). The three transformers: “general”, “specific” and “ultra specific” were trained on the “general”, “specific” and “ultra specific” datasets, respectively. The “ultra specific” dataset (ND12) parameters are a subset of the “specific” dataset (ND11) parameters, which are a subset of the “general” dataset (ND10) parameters. The difference between the accuracy of “general” and “ultra specific” transformers on the “ultra specific” dataset is significant (paired t-test; *p* < 0.05).

### Embedding extraction for downstream tasks

Transformers are composed of two parts, the encoder and the decoder. The encoder creates high dimensional vector representations of the source sentence, i.e., the unaligned sequences, which are passed to the decoder to create the translated sentence, i.e., the aligned sequences. This high-dimensional vector embeds the information in sequences as a numeric representation. We compressed this vector to a vector of a size that does not depend on the number of positions. In the case of *n* sequences, the dimension of the vector is 1024 × (2*n* − 1) (see Methods). To exemplify the utility of such a representation, we used this vector representation as input for a different machine-learning task, which is to estimate the ancestral sequences length that generated the resulting sequences. To this end, we trained a linear regression model that takes the coordinates of the compressed high-dimensional vector as input. The training set includes 90,000 nucleotide MSAs, each with five sequences (ND10). The accuracy of the linear-regression model using these features was evaluated on test data comprising 10,000 MSAs (Fig. 6). The significant correlation between the true and inferred root lengths (*R*^2^ = 0.91 and 2.003 base pairs Mean Squared Error (MSE)), suggests that our approach can be used to compactly code sequences, as a preliminary step for downstream machine-learning tasks.

**Fig. 6.**
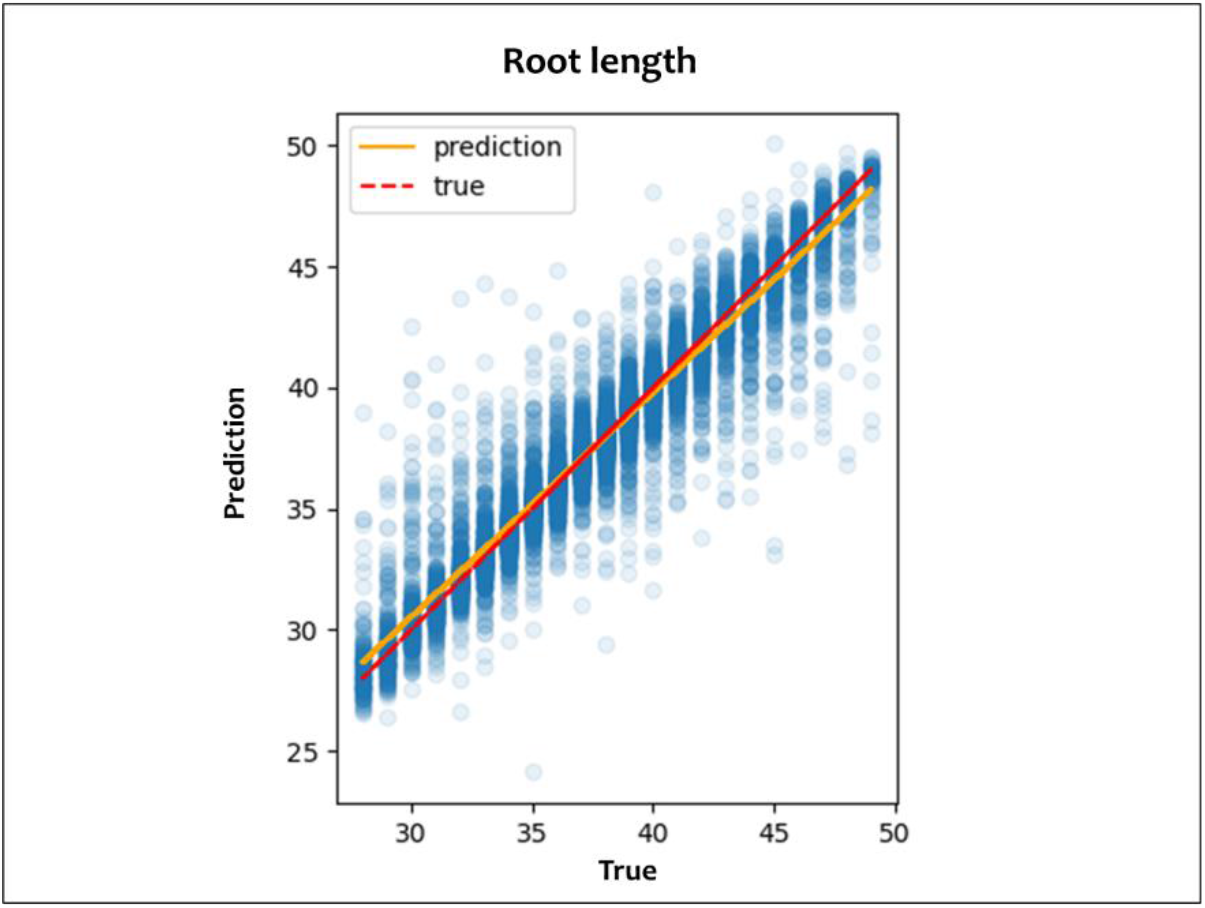
Results of the linear regressor trained to predict the root length from the embedding of the unaligned sequences, with an *R*2 of 0.91 and *MSE* of 2.003 base pairs. The orange line is the regression line, and the red line reflects the *Y* = *x* function. The embeddings are of the ND10 dataset sequences.

### Transfer learning

Our approach heavily depends on transfer learning. Except for the first transformers, for which the weights were randomly initialized, all other transformers used initial weights that were optimized on a previous dataset. The transformers of the nucleotide datasets have a different path of training from the transformers of the amino-acid datasets. In addition, each transformer is optimized based on the previous transformer with the same configuration (as we trained two different transformers for each dataset). To evaluate the contribution of transfer learning to performance, we tested three alternative scenarios (Fig. 7a, see Methods). Briefly, the transformer in scenario 1 (transformer 1) is trained once on a target dataset. Transformer 2 started from the end point of transformer 1 and was retrained on the same target dataset. Transformer 3 (scenario 3) started from the end point of transformer 1 and was trained on various other datasets, and then retrained on the same target dataset. Our results demonstrated the benefit of transfer learning (Fig. 7b). Transformer 3 outperformed transformer 1, both for protein and DNA sequences, with error reduction of 37.3% and 33.3%, respectively (paired t-test; *p* < 0.005). It may be that the increased accuracy resulted from the fact that transformer 3 was trained twice on the target dataset and not due to the additional training. To test this hypothesis, we compared it to transformer 2. Our analysis suggests that some of the improved accuracy is indeed due to the extra training (comparing transformers 1 and 2). Nevertheless, it also shows that transfer learning substantially contributes to performance (comparing transformers 2 and 3), resulting in 16% and 25% error reductions for protein and DNA, respectively (paired t-test; *p* < 0.005).

**Fig. 7.**
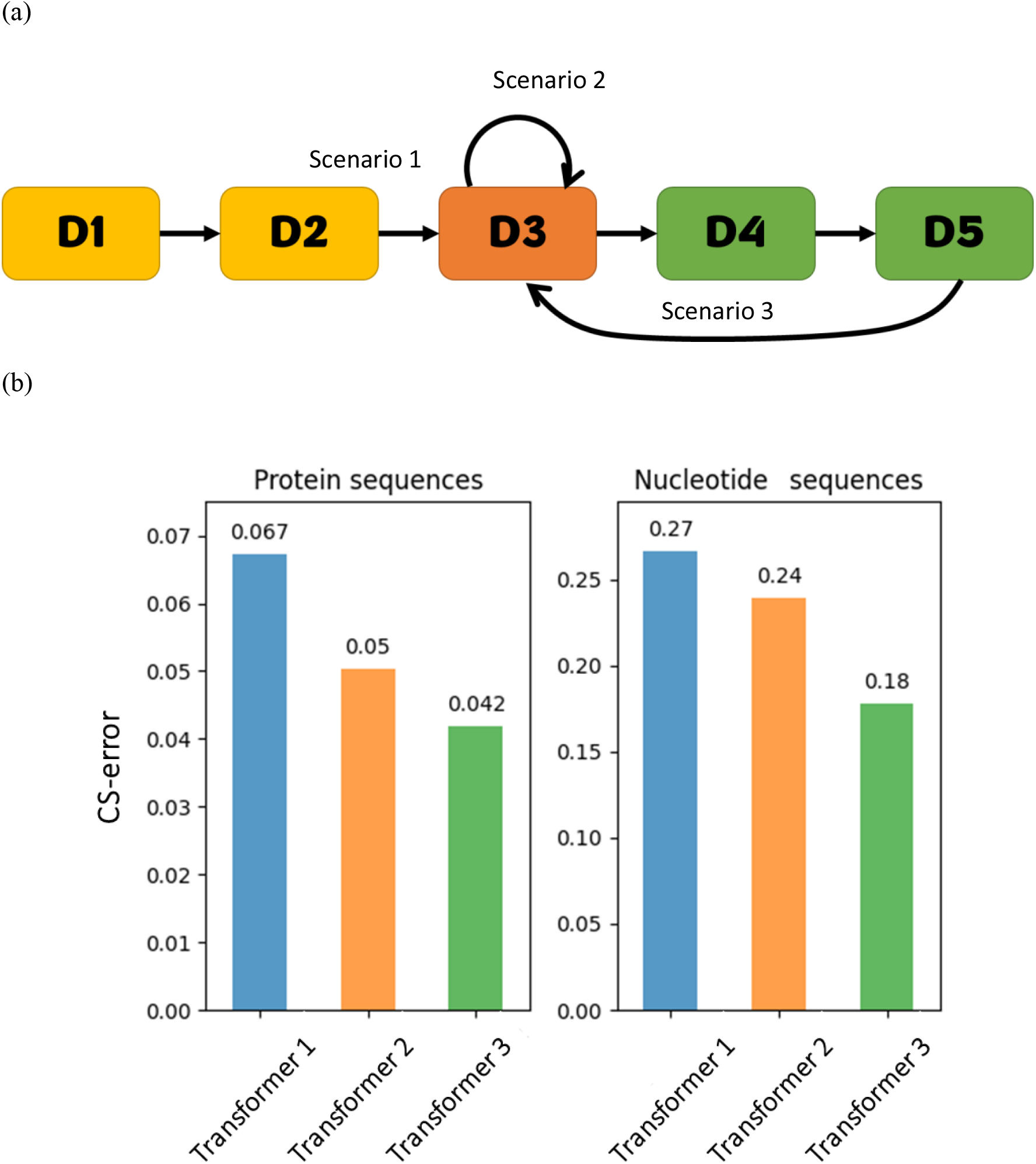
Quantifying the contribution of transfer learning to performance. (a) The transfer learning path. Scenario 1 includes training on “D1”, “D2” and “D3”. Scenario 2 is the same as Scenario 1, but the transformer was trained twice on “D3”. Scenario 3 includes training on “D1”, “D2”, “D3”, “D4”, “D5” and then again on “D3”. “D1” and “D2” represent simpler datasets. “D3” is the target dataset, composed of MSAs of three DNA or amino-acid sequences, on which the performance was evaluated. “D4” and “D5” represent more complex datasets. Arrows between datasets represent the transfer learning path, i.e., the transformer optimized on a dataset was used as a base transformer for the next dataset; (b) The effect of transfer learning on the performance.

### Correlation of certainty and the alignment accuracy

We found a strong dependence between the alignment certainty and the CS-score (Fig. 8). As the certainty of alignments can be calculated by creating multiple alternative alignments for the same set of unaligned sequences (see methods), we could utilize this dependence to infer the most accurate alignment, similar to a previous approach (Edgar, 2022).

**Fig. 8.**
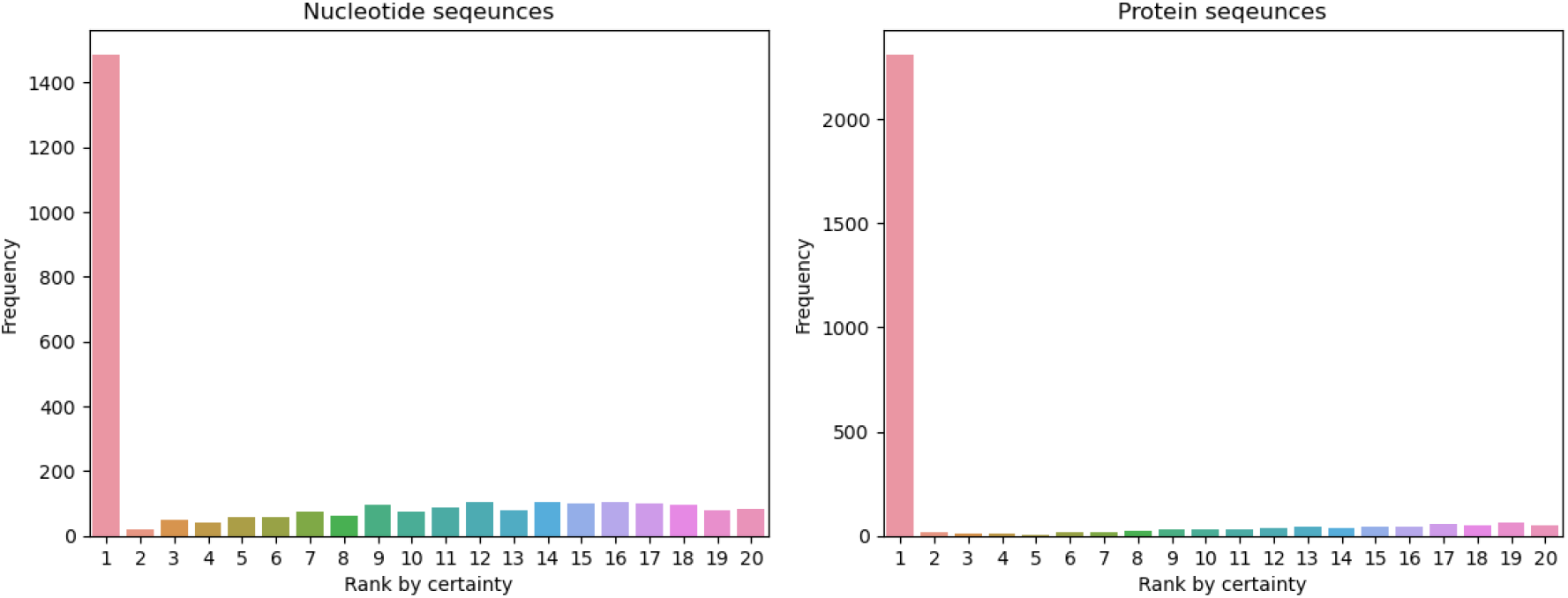
The frequency of the optimal alternative alignment for each certainty rank. For each data point, a total of 20 alternative alignments were considered, each with 10 sequences (SND1 and SPD1 for the nucleotide and protein datasets, respectively). The 20 MSAs were ranked according to their certainty. Next, the most accurate MSA was detected (based on the CS accuracy score) and its ranked recorded. Of note, some of the alternative MSAs may be identical. In case the most accurate MSA was ranked multiple times (e.g., the first and second ranks), we consider its ranked to be the higher rank (e.g., the first). Shown is the distribution of ranks among 3,000 independent data points. A uniform distribution is expected if the certainty rank does not provide any information regarding the alignment accuracy.

Having observed that the alignment with the highest (alignment) certainty is ranked higher than expected (among the set of alternative alignments from a specific dataset), we next directly compared performance between choosing the alignment alternative with the highest certainty and selecting the first alternative alignment. We tested this approach on 10-sequences data points (SND1 and SPD1) and observed a significant CS-error reduction of 9.8% and 20.9% for DNA and protein alignments, respectively (paired t-test; *p* = 0.002).

### Comparing performance

We compared the performance BetaAlign after selecting the MSA with the highest certainty against other commonly used alignment programs, both for DNA and protein sequences (Fig. 9). For DNA sequences, regardless of the number of sequences analyzed, BetaAlign was the most accurate (paired t-test; *p* < 10^−7^), with a minimal error reduction of 12.7%. The second most accurate alignment program was MUSCLE for 4-7 sequences and PRANK for 8-10 sequences. For 10 sequences, for example, BetaAlign had an 13.7% error reduction compared to PRANK (paired t-test; *p* < 10^−12^) and similar results were obtained for other number of sequences. MAFFT, DIALIGN, and ClustalW had a significantly lower performance, with MAFFT outperforming the two other alignment programs. Notably, for protein sequences, BetaAlign was typically the second most accurate. For 10 sequences, the error reduction of PRANK was 5.1% relative to BetaAlign.

**Fig. 9.**
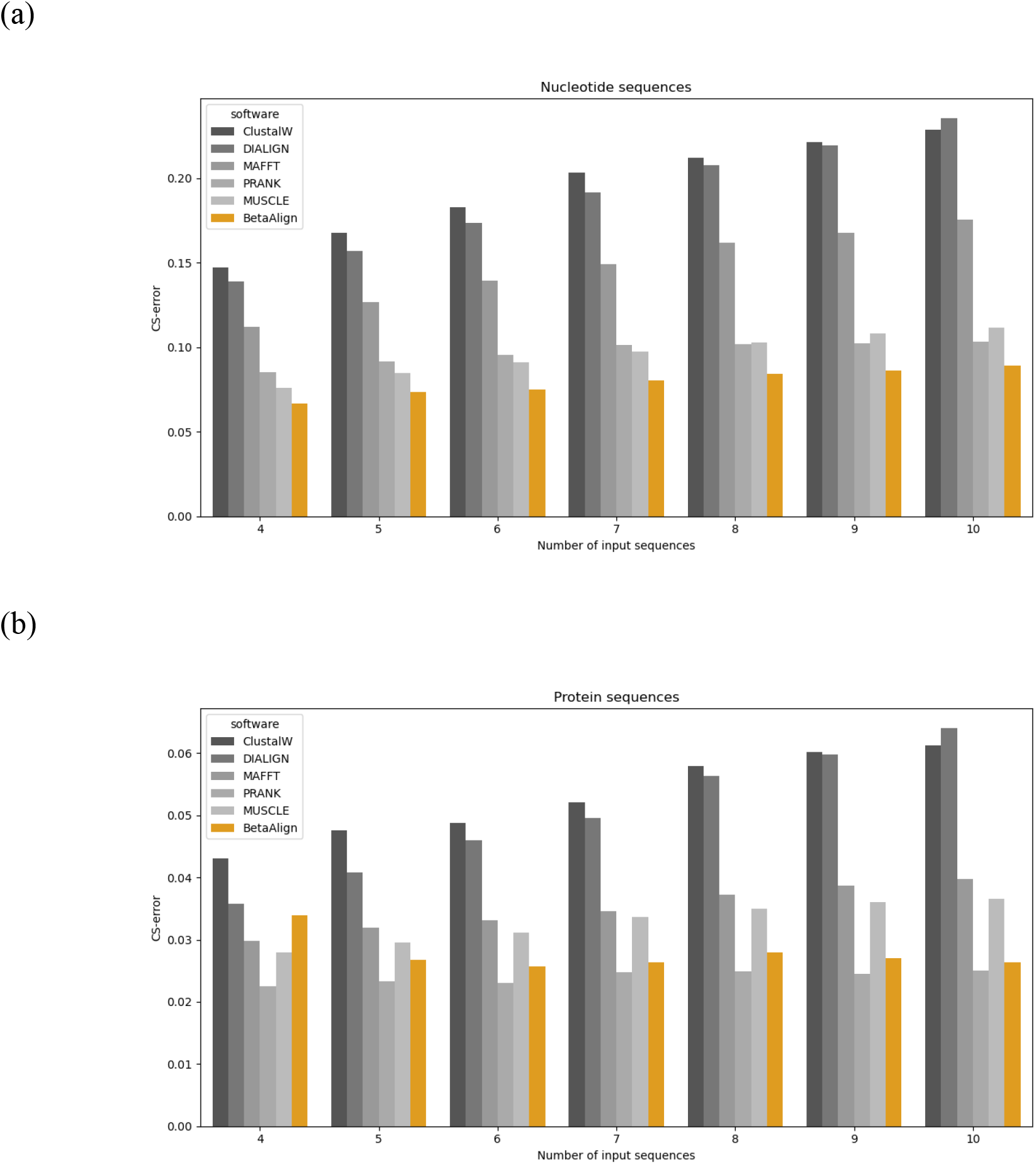
Comparing the results of BetaAlign to different aligners on SND1 (panel (a)) and SPD1 (panel (b)). The y-axis represents the performance of the sequence alignment programs. The lower the CS-error the better the performance.

## Methods

### The generation of training and test data

We first describe in detail the simulation of nucleotide dataset SND1, in which each data point includes ten unaligned sequences and their corresponding ‘true’ MSA. We generated 395,000 and 3,000 data points for training and testing data, respectively. For each data point, we sampled a random tree using the program ETE 3 (Huerta-Cepas et al., 2016), with tree lengths uniformly distributed in the range (0.05, 0.1). The sequences along each tree were simulated using SpartaABC (Loewenthal et al., 2021). Specifically, indel parameters were sampled from the following ranges: *R*_*I*_, *R*_*D*_ ∈ (0.0, 0.05), *A*_*I*_, *A*_*D*_ ∈ (1.01, 2.0), and root length ∈ [32, 44]. Of note, the insertion (*R*_*I*_and *A*_*I*_) and deletion rates (*R*_*D*_and *A*_*D*_) were sampled independently allowing a rich-indel model, in which insertions and deletions can have different evolutionary dynamics. The above parameter ranges were found to accurately describe the indel evolution rates along the tree of life (Loewenthal et al., 2021). The WAG+G and the GTR+G substitution models were used for the protein and nucleotide datasets, respectively. The GTR+G frequencies were (0.37, 0.166, 0.307, 0.158) for the “T”, “C”, “A” and “G”, respectively. Substitution rates were (0.444, 0.0843, 0.116, 0.107, 0.00027) for the “a”, “b”, “c”, “d”, and “e” rate parameters as defined in Yang (1994). These frequencies and rate parameters reflect those that characterize the Yeast Intron Database (Lopez & Séraphin, 2000). Specific information for the simulation of each dataset is provided in Table S2. The datasets are available on HuggingFace (Wolf et al., 2020) at: https://huggingface.co/dotan1111.

### The transformer architecture

We applied the “vaswani_wmt_en_de_big” architecture (Vaswani et al., 2017) with 16 attention heads, embeddings size of 1,024 and 6 layers. We also conducted an experiment to evaluate the effect alternative architectures on performance (see Supplementary Information). We considered a variety of training hyper-parameters configurations for the transformer, including different max tokens values, learning rates, and warmup updates and evaluated them on datasets of pairwise alignments (Supplementary Table S1). We continued to train two configurations that yielded the best results, which we denote as “original” and “alternative”. The max token parameter values were 4,096 and 2,048 for the original and alternative transformers, respectively. For both configurations we used the same learning rate (5E-5) and warmup updates (3,000). Model training and evaluations were executed on a Tesla V100-SXM2-32GB GPU machine.

### Using alternative alignments to increase the accuracy of BetaAlign

A “column certainty” metric was employed to compute “alignment certainty”. Given an alignment, *x*, and a set of alternative alignments, *Y*, the column certainty of each column in *x*, is the number of times the column appears in each alternative alignment *y* ∈ *Y* divided by the total number of alignments in *Y*. As a result, column certainty values range between 0 and 1, where a score of 1 indicates high certainty. The alignment certainty is defined as the average of the column certainty values (Fig. 10).

**Fig. 10.**
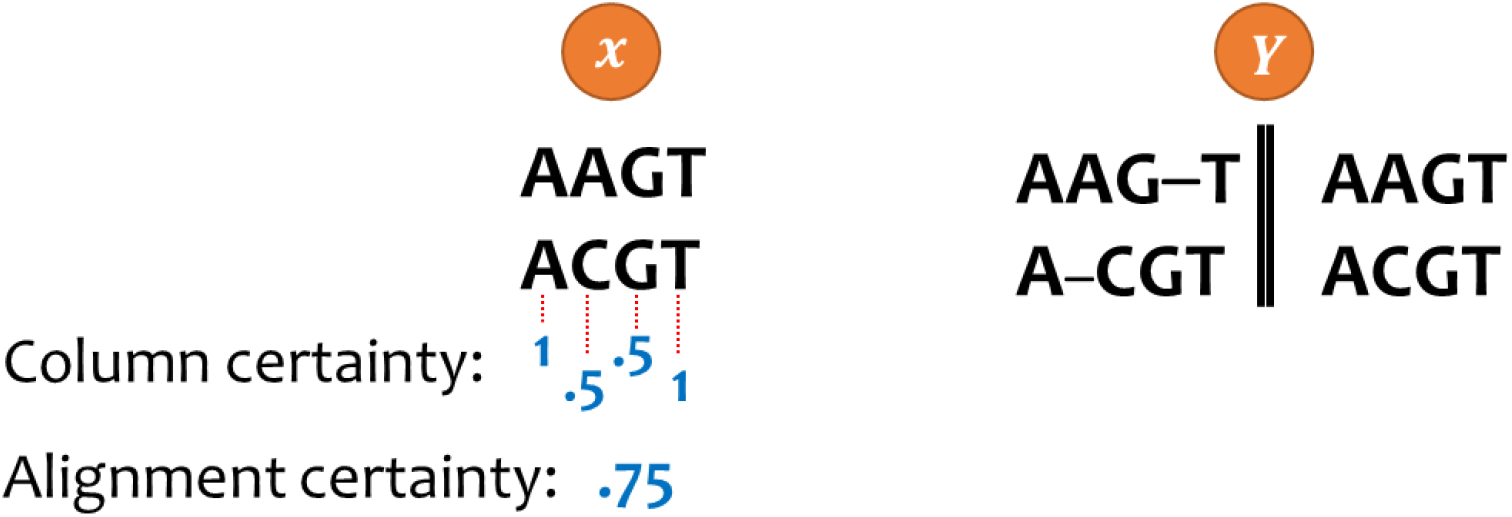
An illustration of calculating the alignment certainty on pairwise alignment. Consider x to be a pairwise alignment where “AAGT” is aligned to “ACGT” and Y to be the collection of two alternative alignments: (1) where “AAG-T” is aligned to “A-CGT” and (2) where “AAGT” is aligned to “ACGT”. To determine the certainty for each column in x, we count the number of appearances in the set of alternative alignments Y and divide it by the size of the set Y. For example, the first column, “AA”, appears both in alignments (1) and (2) and thus its certainty is 2 / 2. The second column, “AC” appears only in alignment (2) and thus its certainty is 1 / 2.

It is possible to generate alternative MSAs for the same set of sequences. For example, alternative MSAs are generated by GUIDANCE to quantify the reliability of different regions within an MSA (Sela et al., 2015). These alternative MSAs are computed by considering alternative guide trees, considering co-optimal solution of pairwise alignments, and changing the alignment scoring scheme. Alternative MSAs are also computed within the alignment program Muscle (Edgar, 2022). The alignment that agrees best with the set of alternative MSAs is then chosen as the inferred MSA. We developed a similar approach for generating alternative MSAs, which is based on the deep learning methodology proposed here. Specifically, we alternate the order of the unaligned sequences given as input to the “*concat*” representation. This results in the inference of different MSAs for the same input. For example, an MSA of three sequences results in six different permutations, thus providing six alternative MSAs and similarly *k*! alternative alignments for *k* sequences. In addition, as we trained several transformers with different training parameters for each dataset, we can add alternative alignments from two or more transformers by processing the same input using these different transformers (Dotan, et al., 2023a).

Formally, let *x*, and ℎ be a list of unaligned sequences, and an aligner program, respectively. When computing alignment, the aligner is dependent on a set of parameters, i.e., a configuration, denoted by *α*. Altering *α* would output a different alignment for the same *x* and ℎ. Thus, for a list of *n* different configurations: *a*_1_,…, *a*_*i*_,…, *a*_*n*_, one would receive *n* different alignments: *Y* = {ℎ_*a*1_ (*x*),…, ℎ_*ai*_(*x*),…, ℎ_*an*_(*x*)}. Of note, the alignments of different configurations could be the same. Creating the different configurations could be done by changing the scoring scheme for the aligners or by changing the permutation of the unaligned sequences in the case of BetaAlign (see above). For each alignment ℎ_*ai*_(*x*) we calculate the alignment certainty described above, by comparing it to all the other alignments *Y* \ {ℎ_*ai*_(*x*)}. We return the alignment that maximizes the alignment certainty. Specifically, we have two transformer configuration (“original” and “alternative”) and for each, we generated 10 alternative MSAs. We return the valid alignment with the highest certainty.

### Evaluating accuracy and coverage

We evaluated the performance of BetaAlign using two metrics: (1) column score (CS), which identifies how many columns are shared between the inferred and the true alignment. Of note, a shared column requires the same characters with the same positions of each character (Sela et al., 2015). The CS is the number of shared columns divided by the number of columns and thus the score is in the range [0,1]. The CS-error is the complementary of the CS to 1; (2) We use the term coverage to denote the percentage of valid alignments from the total number of MSAs generated by the transformer. Examples of invalid alignments are illustrated in Fig. 2.

### Evaluating the effect of training time and size

We generated datasets containing 50,000, 100,000, and 200,000 alignments. Next, we trained transformers on each of the datasets for 60 epochs with the original transformer training parameters. We evaluated the performance of the transformers at the end of each epoch, with respect to the following metrics: (1) training loss, (2) validation loss, (3) fraction of invalid alignments (i.e., 1 – coverage), and (4) CS-error. The validation data contained 2,000 alignments (used to measure the validation loss), and the test data contained 3,000 alignments (used to measure the fraction of invalid alignments and CS-error). Of note, in each of the three experiments we initialized the model with random weights, and thus, transfer learning did not affect these results.

### Evaluating the effect of indel parameters on alignment inference accuracy

To quantify the effect of the evolutionary parameters on alignment inference accuracy we generated training and test data using the same random topology and branch lengths as were used in PD14 (see Table S2). The range of indel evolutionary parameters was binned: For *A*_*I*_ and *A*_*D*_ that dictate indel-length distribution for insertions and deletions, respectively, the following ten bins were considered for each parameter: (1.0, 1.1), (1.1, 1.2)… (1.9, 2.0). For *R*_*I*_and *R*_*D*_that dictate indel rates relative to substitutions for insertions and deletions, respectively, the following ten bins were considered for each parameter: (0.000, 0.005), (0.005, 0.01)… (0.045, 0.05). We thus considered 100 bins for the pair (*A*_*I*_, *A*_*D*_) and similarly for the pair (*R*_*I*_, *R*_*D*_). When analyzing the effect of *A*_*I*_ and *A*_*D*_, for each of the 100 (*A*_*I*_, *A*_*D*_) bins, 100 alignments were generated, in which the *R*_*I*_ and *R*_*D*_ values were sampled randomly from the range (0.00, 0.05). Thus, in total 10,000 MSAs were considered when studying the effect of the *A*_*I*_and *A*_*D*_parameters. Similarly, 10,000 MSAs were considered when studying the effect of the *R*_*I*_and *R*_*D*_parameters, and in this case, in each MSA the *A*_*I*_ and *A*_*D*_ parameters were sampled from the range (1.0, 2.0). The score of the 100 alignments in each bin was averaged to create a total score for each bin.

### Subspace learning evaluation

The MSA in the training data for BetaAlign is generated by evolving sequences along a specific phylogenetic tree and different MSAs are generated with different trees and with different evolutionary models. The substitution and indel dynamics are dictated in this simulation by an evolutionary model (a continuous-time Markov process). Let *g* be the set of evolutionary models and trees used to generate the data. Clearly, a trained aligner, *h*, depends on *g*. In other words, our aligner learns to align sequences generated by the set of evolutionary models *g* that generated the training data. Thus, we can readily create aligners that will best suit a specific subspace of model parameters and trees, e.g., aligners for a specific phylogenetic tree, and similarly aligners for species or proteins with a specific indel or substitution dynamics. In subspace learning, the transformer is optimized on a subspace of the alignment parameters space. To test how subspace learning affects performance, we generated three nucleotide datasets, each one with a narrower range of model parameters, i.e., *A*_*I*_, *A*_*D*_, *R*_*D*_ and *R*_*I*_, branch lengths and root lengths (ND10, ND11, and ND12). We trained BetaAlign starting with the dataset of the widest parameter range (ND10), which we named “general”. Then, the optimized transformers were used as the starting point for additional training on the next dataset, ND11, whose model parameters are a subset of those of ND10. We named this dataset “specific”. The optimized transformers from ND11 were then further trained on the next dataset (ND12) “ultra specific”. Each of the three transformers was evaluated on each of the three test datasets.

### Embedding of MSAs in a high-dimensional space

The deep learning approach presented here enables embedding the information within the sequences in a high-dimensional space, i.e., it allows automatic features extraction, which could be utilized for downstream analyses. The high-dimensional vector is created within the encoding process from a set of unaligned sequences. To obtain the embedded vector, the unaligned sequences were given as an input to the trained transformer. The vector is internally created by the encoder part of the transformer, and we have modified the code of the transformer to extract it (to reduce running time, we skipped the decoder step). This high-dimensional vector contains ∼ 1,024 × *n* × *l* entries, where *n* is the number of input sequences and *l* is the average length of unaligned sequences. A representation of this vector, for three sequences is given in Fig. 11a.

**Fig. 11.**
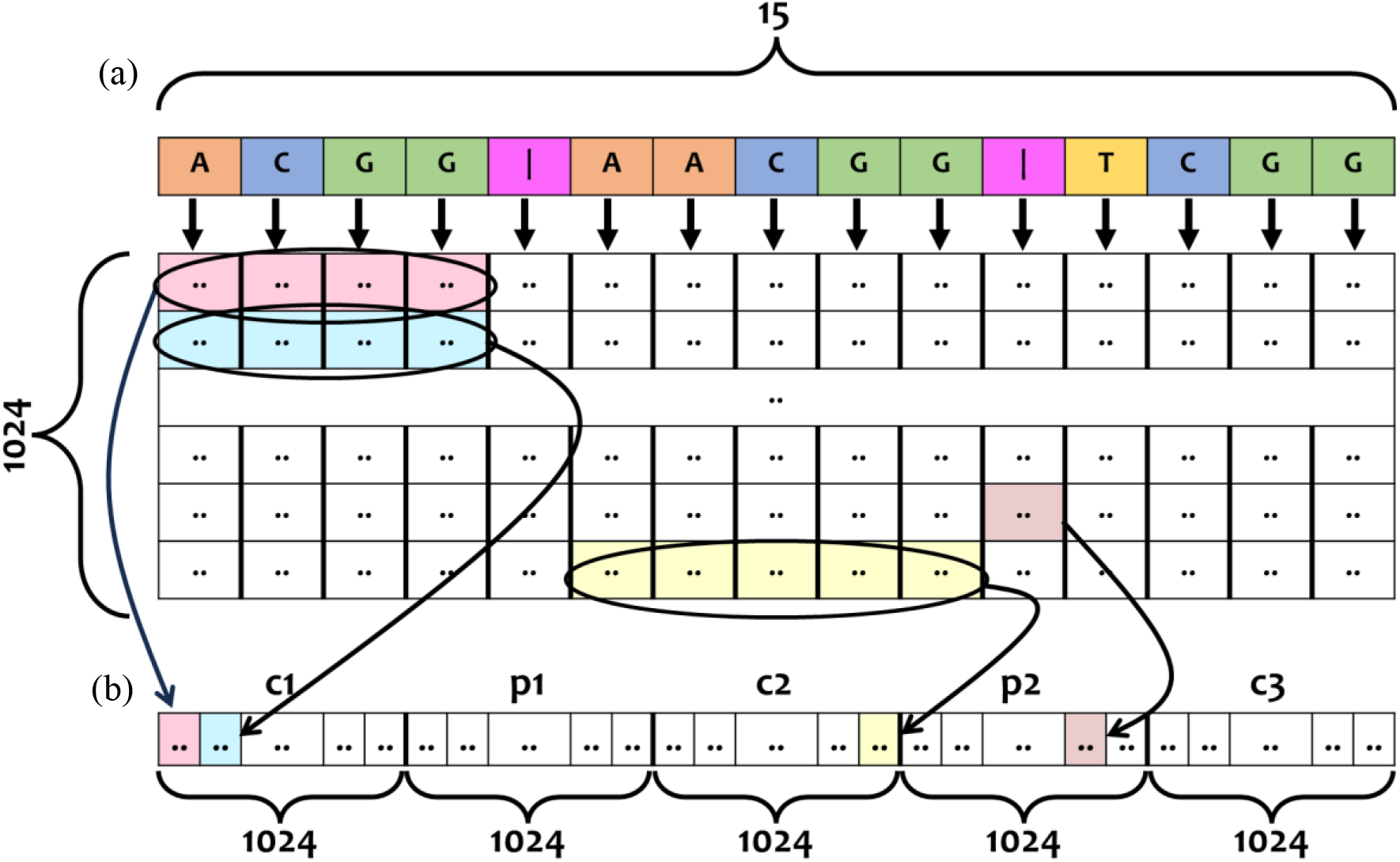
Example of compressing the embedding vector to a fixed size. Consider (a) to be the embedding of three sequences, two of length 4 nucleotides and one of length 5 nucleotides. The embedding dimension is of 1,024 × 15 as there are 15 characters is the input sentence (13 nucleotides and 2 separation characters) and each character is encoded in a numeric vector of size 1,024. The compressed vector is (b) is of size 1 × 5,120 as each one of the input sequences and the pipe sign corresponds to 1,024 entries in the compressed vector. The four first columns in this matrix are averaged and the resulting column vector represents a fixed size vector for the first sequence. This vector is transposed to form c1. The vector representing the pipe character remains the same, it is just transposed to form the vector p1. The next five columns are averaged and transposed to form c2, etc. One should emphasize that the pre-compressed representation already integrates information from all sequences due to the transformer self-attention mechanism, and consequently, the compressed representation also integrates information from all sequences.

For various downstream tasks, it is often desirable to compress this vector to a fixed size, i.e., a size that does not depend on the sequence length (the compressed vector size does depend on the number of sequences). For the compression example shown in Fig. 11, the uncompressed vector is of size 1,024 x 15 and the size of the compressed vector is 1,024 x 5. Each of the unaligned sequences is represented by 1,024 entries in the compressed vector by row-wise averaging of the corresponding tokens in the input sequences. In addition, we use the representations of the pipe character in the compressed vector. Thus, the compressed vector corresponds to a fixed size vector of 1,024 × (2*n* – 1).

### Evaluating and implementing transfer learning

In our work, transfer learning was repeatedly used for training the transformers. The first protein transformer was trained on a simple dataset of pairwise amino acid sequences (we denote this dataset PD1, for protein dataset 1). Its weights were randomly sampled with default values of the Fairseq library (Ott et al., 2019). The resulting trained transformer is termed “PT1”, for protein transformer 1. PT1 was next trained on PD2, resulting in PT2, etc. The term transfer learning is used to denote the fact that in order to obtain PT2, the transformer trained on PD2, was initialized with weights transferred from PT1, rather than random initialization. A similar process was used to train the nucleotide-based transformers (NT1, NT2, etc.) on nucleotide datasets (ND1, ND2, etc.). Of note, transfer learning was applied across this study only between models that processed data with the same representation, i.e., they share the same dictionaries.

We aimed to evaluate the contribution of transfer learning. To this end, we compared three different scenarios (illustrated in Fig. 7). In scenario 1, we evaluate a transformer that first encounters protein data PD5 (three protein sequences). This transformer was trained before on simpler datasets. In scenario 2, the trained transformer from scenario 1 was retrained on PD5, without experiencing more complex datasets. In scenario 3, the trained transformer from scenario 1 was trained on additional more complex datasets (PD6, PD7, PD8, PD9, PD10, PD11, PD12, PD13, PD14, PD15) and was then re-trained on PD5.

A similar evaluation was done on nucleotide transformers. Here instead of PD5, the base-dataset was ND4, comprised of alignments of three sequences. In scenario 3, the additional more complex datasets are: ND5, ND6, ND7, ND8, ND9, ND10, ND11, ND12, ND13, ND14.

### Comparing against other alignment programs

The performance of BetaAlign was compared to the following programs used with default parameters: MUSCLE v3.8.1551 (Edgar, 2004), MAFFT v7.475 (Katoh & Standley, 2013), PRANK v.150803 (Löytynoja & Goldman, 2008), ClustalW 2.1 (Larkin et al., 2007), and DIALIGN dialign2-2 (Morgenstern, 2004).

## Discussion

The weights that are learned by the encoder can be used as a starting point for other machine-learning tasks, i.e., the sequences are embedded as meaningful vectors that hold contextual information. In this work, we demonstrated using such embedding for predicting the length of ancestral sequences, without computing the MSA. A similar approach can be used for other machine-learning tasks, e.g., secondary structure prediction, predicting the stability of proteins, and ancestral sequence reconstruction. In NLP, transferring representations from one task to another is highly common, and encoder-decoder models are commonly used for this purpose (McCann et al., 2017).

There are limitations when using NLP approaches for sequence alignment, one of which arises from the maximum sequence length that can be inserted into an attention-based model. This limitation stems from computing attention matrices, in which the memory requirement increases quadratically with the sequence length. To overcome this issue, we have developed a novel approach that involves splitting and merging the alignment while training the transformer on a slightly different task (Dotan, et al., 2023a). It is possible to apply different techniques to increase the limit on the sizes of the sequences. For example, a different tokenization technique allows multiple amino-acids or nucleotides to be considered as a single token, and thus reduces the number of tokens for the entire sequence (Dotan, et al., 2023b).

We have coupled the NLP domain and the MSA problem by using transformers that were originally designed for natural languages. Thus, future improvements in the NLP field are likely to have a direct impact on future alignment methodologies. We expect that in the next few years, transformers that are dedicated to the task of sequence alignment, together with other breakthroughs in machine learning, will lead to alignment algorithms that account for the specific grammar rules of each set of analyzed sequences and will substantially outperform existing aligners.

## Supplementary Information

### Comparing different transformer architectures

We considered two different architectures for the transformers: “vaswani_wmt_en_de_big” (Vaswani et al., 2017) and “BART” (Lewis et al., 2019). Both types of transformers were trained applying the “*concat*” with the “*spaces*” representations. We tested the results on proteins datasets: PD1, PD2, PD3, and PD4. Of note, for the comparison to be fair, the two transformers were not pre-trained when applied to PD1 and thus their training started from random weights. Both architectures contain 16 attention heads, with an embedding size of 1,024, they differ in the details of their network design, including a different number of layers: 6 and 12 for “vaswani_wmt_en_de_big” and “BART”, respectively. The performance of these architectures was tested with several different sets of internal parameters (max tokens and learning rate). Both the coverage and the CS-score were higher for the “vaswani_wmt_en_de_big” architecture for the two datasets that are most difficult, i.e., PD3 and PD4 (Table S1). We thus selected this architecture for all analyses.

## Supplementary Notes

## Acknowledgments

Edmond J. Safra Center for Bioinformatics at Tel Aviv University Fellowship (ED, EW, NE, MA). TP’s research is supported in part by the Edouard Seroussi Chair for Protein Nanobiotechnology, Tel Aviv University.

---

**Table S1.** We trained two transformer architectures on the same amino-acid datasets and measured the alignment accuracy and coverage. The “*concat*” and “*spaces*” representations were used for input and output transformation, respectively. Datasets PD1, PD2, and PD3 are of pairwise alignments and dataset PD4 includes alignments of three sequences.

| PD1 |  |  |  |  |  |
| --- | --- | --- | --- | --- | --- |
| Transformer name | Architecture | Max tokens | Learning rate | CS-score | Coverage |
| original | BART | 4096 | 5.00E-05 | 0.9996 | **0.9716** |
| alternative | BART | 2048 | 5.00E-05 | **0.9997** | 0.7724 |
| alternative_2 | BART | 4096 | 1.70E-05 | 0.9994 | 0.6161 |
| alternative_3 | BART | 2048 | 1.70E-05 | 0.9995 | 0.6039 |
| original | vaswani_wmt_en_de_big | 4096 | 5.00E-05 | 0.9995 | 0.9508 |
| alternative | vaswani_wmt_en_de_big | 2048 | 5.00E-05 | **0.9997** | 0.6882 |
| alternative_2 | vaswani_wmt_en_de_big | 4096 | 1.70E-05 | 0.9992 | 0.1616 |
| alternative_3 | vaswani_wmt_en_de_big | 2048 | 1.70E-05 | 0.9991 | 0.1436 |

| PD2 |  |  |  |  |  |
| --- | --- | --- | --- | --- | --- |
| Transformer name | Architecture | Max tokens | Learning rate | CS-score | Coverage |
| original | BART | 4096 | 5.00E-05 | 0.9922 | 0.2279 |
| alternative | BART | 2048 | 5.00E-05 | **0.9983** | 0.8167 |
| original | vaswani_wmt_en_de_big | 4096 | 5.00E-05 | 0.997 | **0.8352** |
| alternative | vaswani_wmt_en_de_big | 2048 | 5.00E-05 | 0.9978 | 0.6348 |

| PD3 |  |  |  |  |  |
| --- | --- | --- | --- | --- | --- |
| Transformer name | Architecture | Max tokens | Learning rate | CS-score | Coverage |
| original | BART | 4096 | 5.00E-05 | 0.7427 | 0.5976 |
| alternative | BART | 2048 | 5.00E-05 | 0.7644 | 0.5576 |
| original | vaswani_wmt_en_de_big | 4096 | 5.00E-05 | 0.8389 | **0.6464** |
| alternative | vaswani_wmt_en_de_big | 2048 | 5.00E-05 | **0.9362** | 0.511 |

| PD4 |  |  |  |  |  |
| --- | --- | --- | --- | --- | --- |
| Transformer name | Architecture | Max tokens | Learning rate | CS-score | Coverage |
| original | BART | 4096 | 5.00E-05 | 0.9785 | 0.5054 |
| alternative | BART | 2048 | 5.00E-05 | 0.9878 | 0.8405 |
| original | vaswani_wmt_en_de_big | 4096 | 5.00E-05 | 0.993 | **0.995** |
| alternative | vaswani_wmt_en_de_big | 2048 | 5.00E-05 | **0.9949** | 0.9794 |

**Table S2.** Sparta ABC indel parameters as follow: for the rate of insertion (*RI*), for the rate of deletion (*RD*), a parameter for the insertion Zipfian distribution (*A*_*I*_), a parameter for the deletion Zipfian distribution (*A*_*D*_) and the root length. Which the latter is sampled uniformly from (*min range* × 0.8, *max range* × 1.1). The order of the datasets refers to the order in which the transformers were trained. For example, the first nucleotide transformer was trained on dataset ND1, then the optimized weights were the starting point of ND2, etc. Tables (a), (b), (c) refer to nucleotide, protein datasets and a special table for specific datasets, respectively. “S” at the start of the dataset name, refers to a special dataset.

| Dataset name | Branch length | Root length | $R_I$ & $R_D$ | $A_I$ & $A_D$ | Number input sequences |
| --- | --- | --- | --- | --- | --- |
| ND1 | 0.03 - 0.1 | 50 - 60 | 0.0 - 0.05 | 1.01 - 2.0 | 2 |
| ND2 | 0.03 - 0.3 | 100 - 300 | 0.0 - 0.05 | 1.01 - 2.0 | 2 |
| ND3 | 0.3 - 0.6 | 200 - 300 | 0.04 - 0.05 | 1.01 - 2.0 | 2 |
| ND4 | 0.1 - 0.3 | 50 - 60 | 0.04 - 0.05 | 1.01 - 2.0 | 3 |
| ND5 | 0.15 | 55 | 0.5 | 1.0 - 1.01 | 3 |
| ND6 | 0.15 | 55 | 0.5 | 1.01 | 3 |
| ND7 | 0.15 | 55 | 0.5 | 1.5 | 3 |
| ND8 | 0.05 - 0.1 | 55 | 0.0 - 0.05 | 1.01 - 2 | 4 |
| ND9 | 0.05 - 0.1 | 55 | 0.03 - 0.05 | 1.01 - 2 | 4 |
| ND10 | 0.07 - 0.1 | 35 - 45 | 0.0 - 0.05 | 1.01 - 2 | 5 |
| ND11 | 0.08 - 0.09 | 37 - 42 | 0.03 - 0.05 | 1.01 - 2 | 5 |
| ND12 | 0.09 | 40 | 0.04 | 1.3 | 5 |
| ND13 | 0.05 - 0.1 | 55 | 0.02 - 0.03 | 1.0 - 1.1 | 4 |
| ND14 | 0.9 | 40 | 0.01 - 0.02 | 1.35 - 1.45 | 5 |
| ND15 | 0.07 - 0.1 | 35 - 45 | 0.0 - 0.05 | 1.01 - 2 | 7 |
| ND16 | 0.07 - 0.1 | 70 - 80 | 0.0 - 0.05 | 1.01 - 2 | 7 |

(b)
| Dataset name | Branch length | Root length | $R_I$ & $R_D$ | $A_I$ & $A_D$ | Number input sequences |
| --- | --- | --- | --- | --- | --- |
| PD1 | 0.03 - 0.05 | 30 - 40 | 0.0 - 0.05 | 1.01 - 2.0 | 2 |
| PD2 | 0.05 - 0.1 | 70 - 80 | 0.04 - 0.05 | 1.01 - 2.0 | 2 |
| PD3 | 0.1 - 0.3 | 200 - 250 | 0.04 - 0.05 | 1.01 - 2.0 | 2 |
| PD4 | 0.03 - 0.1 | 30 - 40 | 0.0 - 0.05 | 1.01 - 2.0 | 3 |
| PD5 | 0.1 - 0.2 | 50 - 60 | 0.04 - 0.05 | 1.01 - 2.0 | 3 |
| PD6 | 0.15 | 50 | 0.05 | 1.01 | 3 |
| PD7 | 0.15 | 50 | 0.05 | 1.5 | 3 |
| PD8 | 0.05 - 0.1 | 30 - 40 | 0.0 - 0.05 | 1.01 - 2 | 4 |
| PD9 | 0.075 | 35 | 0.03 | 1.07 | 4 |
| PD10 | 0.04 - 0.08 | 30 - 40 | 0.0 - 0.05 | 1.01 - 2 | 5 |
| PD11 | 0.04 - 0.08 | 30 - 40 | 0.0 - 0.05 | 1.01 - 2 | 6 |
| PD12 | 0.1 | 30 - 40 | 0.0 - 0.05 | 1.01 - 2 | 6 |
| PD13 | 0.1 | 30 - 40 | 0.03 - 0.05 | 1.01 - 2 | 6 |
| PD14 | 0.07 - 0.1 | 25 - 35 | 0.0 - 0.05 | 1.01 - 2 | 7 |
| **PD15** | 0.08 - 0.09 | 27 - 32 | 0.03 - 0.05 | 1.01 - 2 | 7 |
| **PD16** | 0.09 | 30 | 0.04 | 1.3 | 7 |
| **PD17** | 0.05 - 0.1 | 40 | 0.0 - 0.05 | 1.01 - 2 | 10 |
| **PD18** | 0.07 - 0.1 | 25 - 35 | 0.04 - 0.05 | 1.01 - 2 | 7 |

(c)
| **Dataset name** | **Branch length** | **Root length** | **$R_I$ & $R_D$** | **$A_I$ & $A_D$** | **Number input sequences** |
| --- | --- | --- | --- | --- | --- |
| **SND1** | 0.05 - 0.1 | 40 | 0.0 - 0.05 | 1.01 - 2.0 | 10 |
| **SPD1** | 0.05 - 0.1 | 40 | 0.0 - 0.05 | 1.01 - 2.0 | 10 |
| **SPD2** | 0.05 - 0.1 | 40 | 0.0 - 0.05 | 1.01 - 2 | 10 |
| **SPD3** | 0.07 - 0.1 | 25 - 35 | Dynamic | 7 |  |

